# A screen of FDA-approved drugs identifies inhibitors of Protein Tyrosine Phosphatase 4A3 (PTP4A3 or PRL-3)

**DOI:** 10.1101/2020.05.24.098665

**Authors:** Dylan R. Rivas, Mark Vincent C. Dela Cerna, Caroline N. Smith, Blaine G. Patty, Donghan Lee, Jessica S. Blackburn

## Abstract

Protein tyrosine phosphatase 4A3 (PTP4A3 or PRL-3) is highly expressed in a variety of cancers, where it promotes tumor cell migration and metastasis leading to poor prognosis. Despite its clinical significance, there are currently no viable options to target PRL-3 *in vivo*. Here, we screened 1,443 FDA-approved drugs for their ability to inhibit the activity of the PRL phosphatase family. We identified five specific inhibitors for PRL-3 as well as one selective inhibitor of PRL-2. Additionally, we found nine drugs that broadly and significantly suppressed PRL activity. Two of these broad PRL inhibitors, Salirasib and Candesartan blocked PRL-3-induced migration in human embryonic kidney (HEK) cells with no negative impact on cell viability. Both drugs prevented migration of PRL-3 expressing human colorectal cancer cells to a similar degree as the research-grade PRL inhibitor, Thienopyridone, and are selective towards PRLs over other phosphatases. *In silico* modeling revealed that Salirasib binds a putative allosteric site near the WPD loop of PRL-3, while Candesartan binds a potentially novel targetable site adjacent to the CX_5_R motif. Inhibitor binding at either of these sites is predicted to trap PRL-3 in a closed conformation, preventing substrate binding and inhibiting function.

## Introduction

Phosphatases work in concert with kinases to control phosphorylation levels of proteins, lipids, and other macromolecules to regulate many, if not all, cellular processes. Consequently, dysregulated phosphorylation is a hallmark of cancer and multiple other diseases. While kinases, which catalyze phosphorylation, have been drug targets for decades in cancer research, interest in phosphatase drug discovery is increasing as the critical roles of phosphatases in oncogenesis and cancer progression are beginning to be appreciated. Currently more than thirty potential oncogenic phosphatases have been identified, with roles in cellular proliferation, differentiation, migration, and angiogenesis, among others^1^. Protein tyrosine phosphatases (PTPs), in particular, have emerged as central regulators of cancer development and progression, with their increased activity correlated with enhanced tumor formation in mouse models and with worse prognosis in patients^2–5^.

Members of the phosphatase of regenerating liver (PRL) family, also known as the Protein Tyrosine Phosphatase 4A (PTP4A) family, are dual specificity phosphatases. While their normal cellular functions are largely unknown, the PRL family has been repeatedly shown to be involved in cancer progression. In particular, PRL-3 is a well-defined biomarker of metastasis in multiple cancer types, including melanoma, colorectal, and ovarian cancers, where PRL-3 expression is significantly higher in metastatic lesions compared to the primary tumor site^6–14^. In a comprehensive study of 151 patient samples across eleven common human tumors types, PRL-3 protein expression was upregulated in 80.6% of tumor samples compared to matched normal tissue^15^. High PRL-3 expression is also associated with worse prognosis in human leukemia, breast, gastric, ovarian, and colorectal cancer^8,16–19^. The function of PRL-3 in cancer progression is now well-documented. PRL-3 over-expression in tumors and normal cells demonstrated that it plays roles in inhibiting apoptosis, promoting epithelial to mesenchymal transition (EMT), and inducing migration. Additionally, *in vivo*, PRL-3 over-expression results in accelerated tumor formation and increased metastasis across a variety of tumor types^20–23^. Conversely, PRL-3 loss has been shown to prevent tumor growth and metastasis in several *in vivo* models^24–26^. In one example, PRL-3 loss resulted in 50% less tumor formation in a colitis-associated colorectal cancer model^27^.

The other PRL family members, PRL-1 and PRL-2, share a high degree of sequence homology to PRL-3 and may possess similar functions. Like PRL-3, both PRL-1 and PRL-2 have been shown to prevent contact inhibition, increase tumor growth, and enhance cell migration and invasion^28–32^. Additionally, high PRL-1 and PRL-2 expression has been reported in a variety of cancer types including cervical, hepatic, and breast cancers ^33–35^. Although less studied than PRL-3, data indicate that PRL-1 and PRL-2 overexpression increases metastasis in mouse models, while their loss decreases tumor cell migration and invasion. Together these results demonstrate the importance of the PRL family both in tumor formation and cancer progression, which has made them attractive therapeutic targets.

Currently, there are no clinically available PRL inhibitors. This is in large part due to several significant challenges associated with small molecule inhibition of the PRL family, including the high level of homology between the PRLs, which makes targeting individual PRLs difficult, and conservation of the active site between PRLs and other tumor suppressive PTPs, such as PTEN^36,37^. Additionally, the PRL active site is shallower, wider, and more hydrophobic than other phosphatases, making design of PRL-specific inhibitors more challenging. In spite of these obstacles, several groups have identified or developed PRL-specific inhibitors including Theinopyridone, JMS-053, Compound 43, and Analog 3 ^38–42^. While these compounds exhibit anti-cancer effects *in vitro* and in mouse xenograft studies, pharmacokinetic concerns have thus far prevented further development into clinical and therapeutic agents. These compounds also target all PRLs, raising concerns for off-target effects, since the physiologic functions of the family are unknown. Promisingly, a humanized PRL-3 antibody was recently developed that prevents growth of PRL-3 expressing tumors, while not targeting PRL-3-negative tissues. However, this antibody is not yet in clinical trials and could be a decade or more away from widespread availability.

To address the immediate need for PRL-3 inhibitors in the clinic, we screened a library of 1,443 FDA approved drugs for their ability to modulate the phosphatase activity of PRLs. We found one selective inhibitor of PRL-2, five selective inhibitors of PRL-3, and nine potent inhibitors of the PRL family. Two drugs from the latter group, Salirasib and Candesartan, prevented PRL-3 mediated cell migration in PRL-3 over-expressing human embryonic kidney (HEK) cells without impacting cell viability. Furthermore, these compounds were able to inhibit migration in multiple colorectal cancer cell lines that expressed high levels of PRL-3. *In silico* docking of the drugs to PRL-3 showed that Salirasib binds to PRL-3 in the same site proposed for the research-grade PRL-3 inhibitor, JMS-053, while Candesartan binds a secondary site in PRL-3 that has not previously been targeted. These drugs appear to function allosterically by locking PRLs in the closed confirmation. Together, our results indicate that it may be possible to repurpose FDA-approved drugs to block PRL-3 activity in human cancer cells. These drugs may also provide some insight into the structure of compounds that are best able to selectively target PRL-3 and can be used as chemical probes to uncover biological functions of PRL-3.

## Results

### Identification of FDA-approved drugs that inhibit PRL activity

Phosphatase activities of recombinantly expressed PRL-1, PRL-2, and PRL-3 were evaluated based on their ability to hydrolyze the synthetic substrate, phospho-tyrosine analog 6,8-Difluoro-4-methylumbelliferyl phosphate (DiFMUP). DiFMUP hydrolysis could be significantly inhibited by three research-grade PRL inhibitors: the rhodanine derivative PRL Inhibitor I^43^, Analog 3^40^, and Thienopyridone^39^ (Supplemental Fig. 1). Similar to prior reports, Thienopyridone was most effective, blocking PRL activity by ~95%, and was used as a positive control for PRL inhibition throughout the rest of our studies. We next screened 1,433 FDA-approved drugs for their ability to inhibit PRL dephosphorylation of DiFMUP. Phosphatase activity after drug treatment ranged from 0-150% of the dimethylsulfoxide (DMSO) control. Fifty-three compounds that emit fluorescence autonomously were excluded due to the potential interference with data analysis. The mean phosphatase activity across the remaining 1,380 drugs was calculated for each PRL, and a compound was considered a hit in this assay when the drug reduced the phosphatase activity of the PRL to three standard deviations lower than the mean phosphatase activity (Fig. 1a-b, Supplemental Fig. 2). Supplemental Table 1 shows the initial screening data for each drug against PRL-1, PRL-2, and PRL-3, and those meeting the cut-off are highlighted. The DiFMUP assay was then repeated on the initial hits, which confirmed nine broad inhibitors that block the phosphatase activity of PRL-1, PRL-2, and PRL-3 by more than 80% (Fig. 1c, Supplemental Fig. 3). Additionally, there was a single drug that preferentially inhibited PRL-2, and five drugs that preferentially inhibited PRL-3 over other PRLs (Fig. 1d-e).

**Figure 1.**
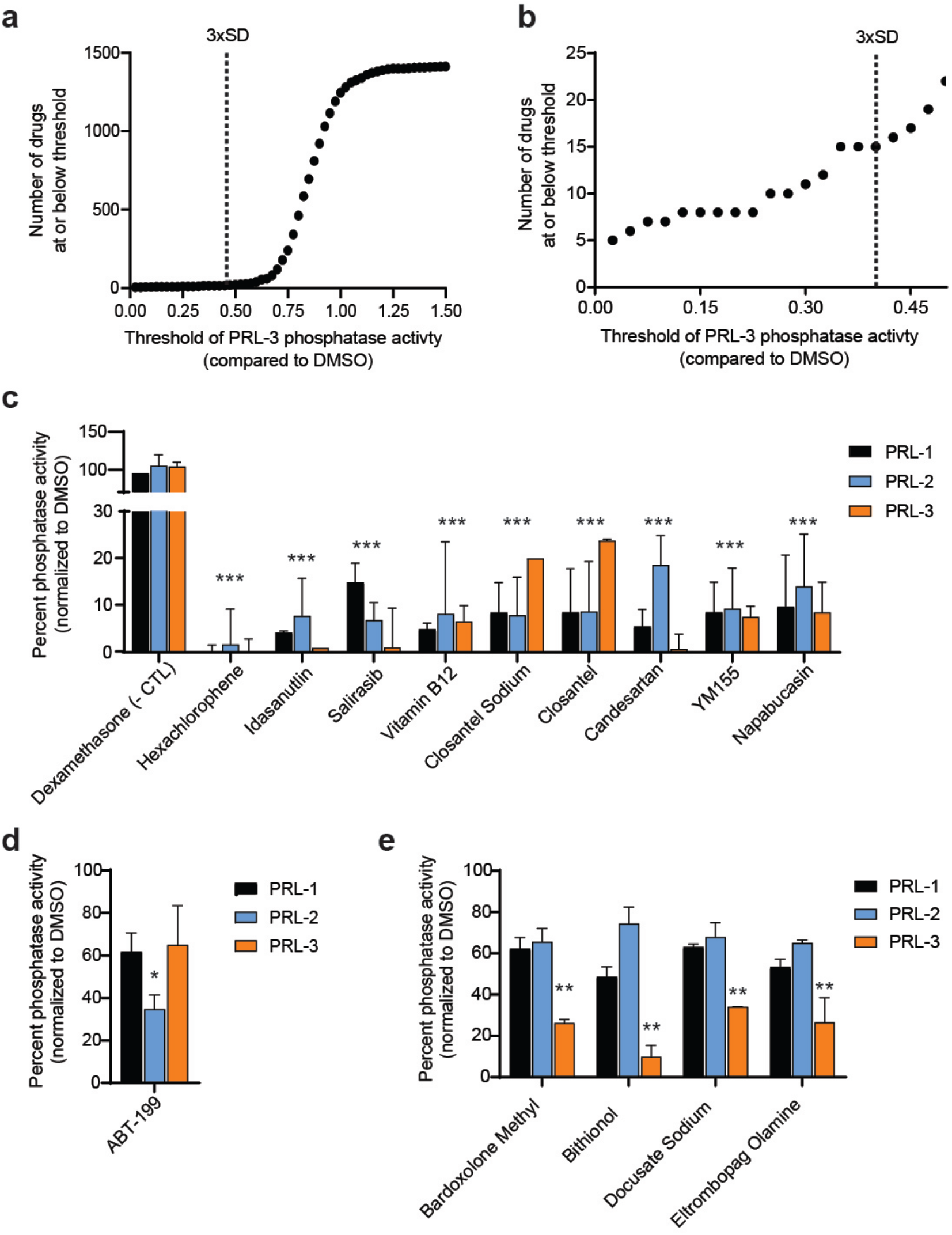
Multiple FDA-approved compounds inhibit the phosphatase activity of PRL proteins. (**a**) S-curve showing the number of drugs at or below the threshold PRL-3 phosphatase activity, compared to DMSO. The PRL-3 phosphatase activity across all drugs was averaged, and three-fold standard deviations (3xSD) below the mean was used to identify hits in the screen. (**b**) Inset of A, showing the number of drugs able to inhibit PRL-3 phosphatase activity 3-fold standard deviations from the mean. (**c**) Positive hit validation for PRL family inhibitors. The FDA-approved drug Dexamethasone, which has no effect on PRL-3 phosphatase activity, was used as a negative control (- CTL). (**d**) ABT-199 demonstrates selectivity in inhibiting PRL-2. (**e**) Hit validation of PRL-3 specific inhibitors. All assay where ran at a final PRL protein concentration of 2.5 μM, a drug concentration of 40 μM, and DiFMUP concentration at the previously reported K_M_ of the protein. The screen of all compounds was run with technical duplicates. Bars represent the average phosphatase activity of the initial screen plus two additional independent experiments. Error bars represent standard deviation. * p < 0.05, **p < 0.001, *** p < 0.0001 by either two way ANOVA with Dunnet’s correction (c) or one way ANOVA with Tukey’s HSD (d,e).

All broad inhibitors blocked PRL phosphatase activity in a dose dependent manner, between 1 nM and 100 μM. The broad PRL inhibitors loosely separated into those with high IC_50_ values (greater than 1 μM), and low IC_50_ values (less than 1 μM) for PRL-1, PRL-2 and PRL-3 (Fig. 2a-b, Supplemental Fig 4). Additionally, the PRL-3 specific inhibitors were also able to selectively inhibit PRL-3 phosphatase activity at a significantly lower IC50 than PRL-1 and PRL-2 (*p* < 0.0001), although they did inhibit these phosphatases to an extent (Fig. 2c-d, Supplemental Fig 5). On average the broad PRL inhibitors had an IC_50_ value 3-fold lower than the specific PRL-3 inhibitors (Table 1), so we focused on the broad PRL inhibitors. The specific PRL-2 inhibitor, ABT-199, is also a potent inhibitor of Bcl-2^44^. Due to the probable associated toxicity at the dose of drug needed for effective PRL-2 inhibition, this drug was not pursued further.

**Figure 2.**
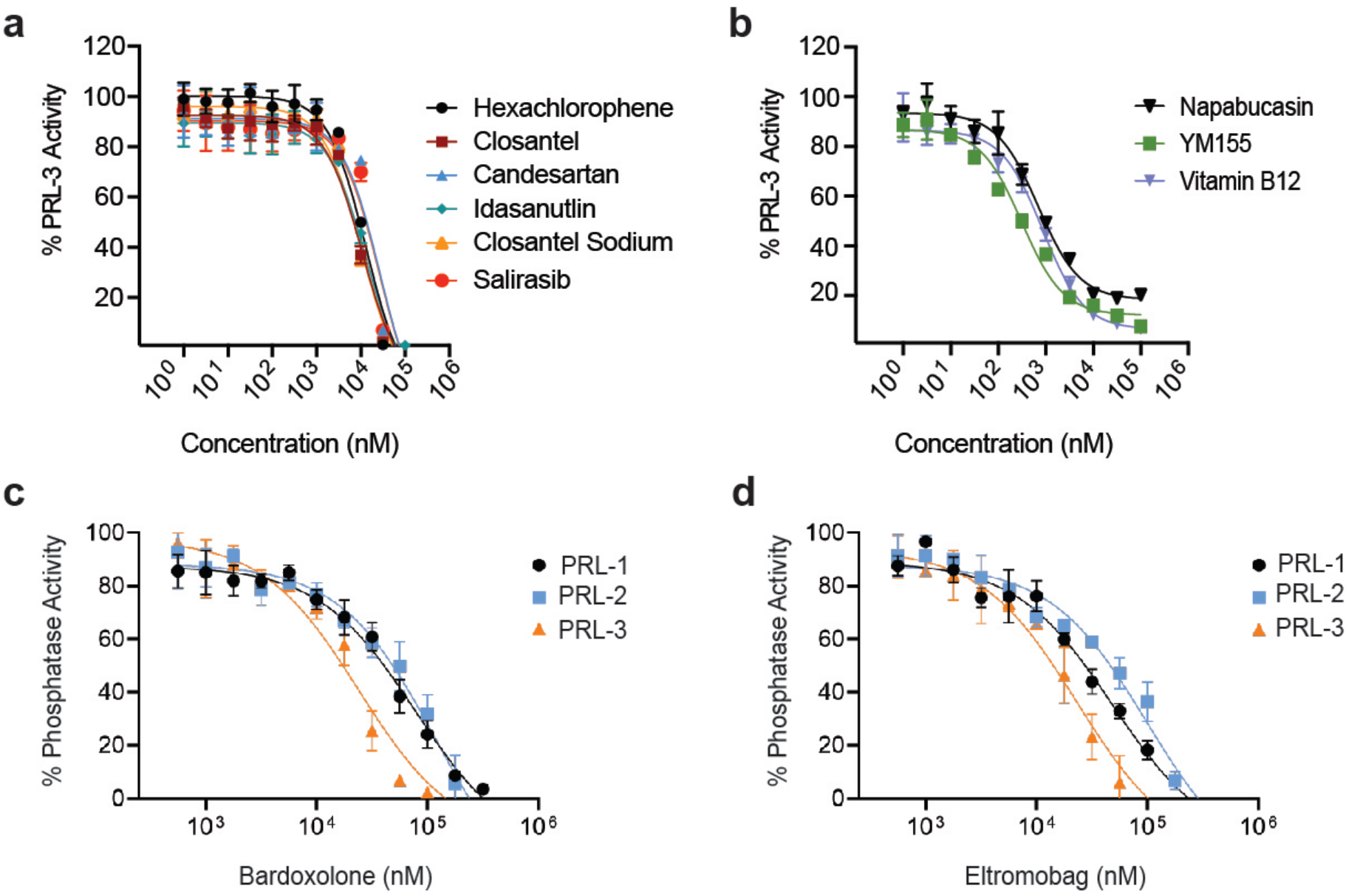
Dose dependent inhibition of PRL family members with FDA-approved drugs. Dose response curves showing DiFMUP phosphatase activity of PRL-3 when treated with doses ranging between1 nM and 100 μM of drug. Inhibition of PRL-3 phosphatase activity by the broad PRL family inhibitors are shown, and drugs are subdivided into those with (**a**) high IC_50_ values and (**b**) low IC_50_ values. A subset of compounds preferentially inhibited PRL-3, including (**c**) Bardoxolone and (**d**) Eltromobag. All assays utilized a final protein concentration of 2.5 μM with DiFMUP concentration at the K_M_ of the protein, and were run in technical duplicates in three independent experiments. Data represent mean phosphatase activity and error bars represent standard deviation between assays.

**Table 1:** Calculated IC_50_ values in μM from broad and specific inhibitors of PRL phosphatase activity

|  | PRL-1 | PRL-2 | PRL-3 |
| --- | --- | --- | --- |
| **Broad Inhibitors** |  |  |  |
| Candesartan | 69 ± 8 | 80 ± 15 | 28 ± 4 |
| Closantel | 17 ± 3 | 14 ± 5 | 11 ± 0.5 |
| Closantel Sodium | 22 ± 8 | 18 ± 1 | 10 ± 0.4 |
| Hexachlorophene | 31 ± 4 | 35 ± 10 | 13 ± 1 |
| Idasanutlin | 20 ± 3 | 16 ± 2 | 13 ± 0.4 |
| Napabucasin | 0.4 ± 0.2 | 1.1 ± 0.3 | 0.7 ± 0.1 |
| Salirasib | 53 ± 5 | 118 ± 22 | 27 ± 3 |
| Vitamin B12 | 0.7 ± 0.2 | 0.9 ± 0.2 | 0.9 ± 0.1 |
| YM155 | 0.3 ± 0.1 | 0.7 ± 0.1 | 0.3 ± 0.1 |
| **PRL-3 Specific Inhibitors** |  |  |  |
| Bardoxolone | 85 ± 43 | 136 ± 75 | 24 ± 5 * |
| Bithionol | 79 ± 25 | 115 ± 34 | 33 ± 2 * |
| Docusate Sodium | 94 ± 26 | 140 ± 44 | 43 ± 5 * |
| Eltromobag | 56 ± 10 | 105 ± 19 | 26 ± 11 * |
| Eltromobag Olamine | 61 ± 15 | 166 ± 22 | 27 ± 4 * |
| Embelin | 114 ± 16 | 124 ± 60 | 52 ± 9 * |

\*p < 0.05 between IC<sub>50</sub> values for PRL-3 and PRL-1 and PRL-2 by one-way ANOVA with Tukey's HSD.

### A subset of compounds is non-toxic to a human cell line at doses effective at inhibiting PRL-3

Drugs were next tested for off-target effects on normal cell viability at the IC_50_ of PRL-3 inhibition, using the immortalized human embryonic kidney cell line 293T (HEK293T). Thienopyridone, Vitamin B12, and Salirasib, do not effect on viability at drug doses ranging from 1-100 μM (Fig. 3a), measured by 3-(4-5-dimethylthiazol-2-yl)-2,5-diphenyltetrazolium bromide (MTT) absorbance after 16 hours of incubation with the compounds. Candesartan showed insignificant effect on viability up to the concentration of 50 μM but viability dropped significantly at 100 μM, indicating potential toxicity of the compound at concentrations well above the IC_50_ for PRL-3 inhibition. Hexachlorophene and Closantel increased MTT absorbance in 293T cells (Fig. 3b); this result likely indicates an increase in cellular proliferation and cell number, rather than an increase in viability. Finally, Idasanutlin, YM155, and Napabucasin decreased viability of 293T cells (Fig. 3c), most likely due to these drugs having a known higher affinity for proteins involved in apoptotic pathways^45,46^ and potentially other off-target effects. As the ultimate goal of this study is to identify drugs that could be used as anti-cancer therapy in PRL-3 expressing tumors, we eliminated drugs that enhanced cell proliferation, and focused on those that decreased or caused no change in cell viability.

**Figure 3.**
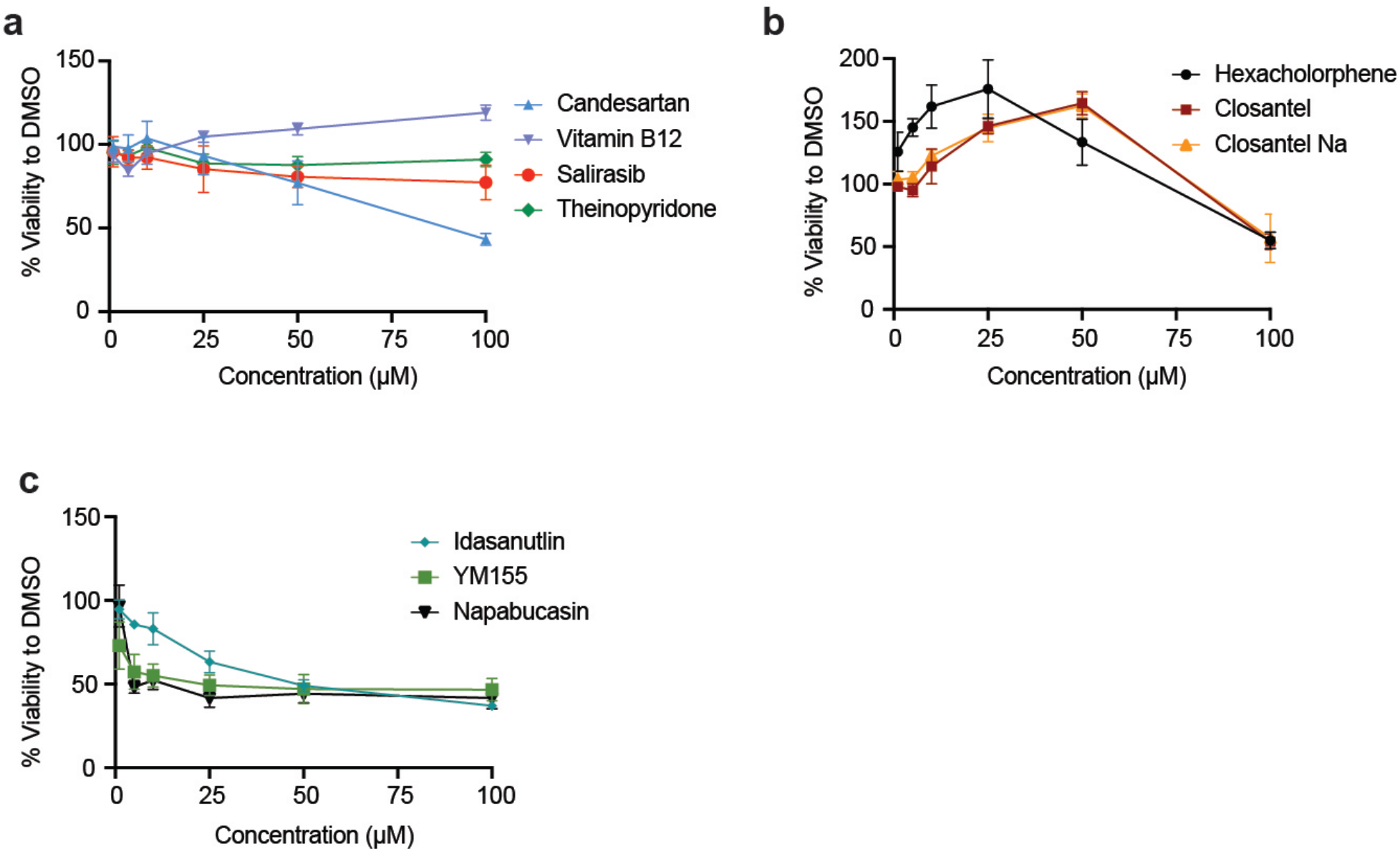
Salirasib and Candesartan are non-toxic PRL inhibitors. HEK293T cells were cultured with drugs at a range of concentrations and viability assessed after 16 hours. Drugs were sub-divided into those that were (**a**) minimally toxic, (**b**) caused increased viability, and (**c**) were highly toxic. Viability was measured as cellular reduction potential using MTT dye, and assays were run in triplicate in three independent experiments. Data represent mean viability and error bars represent standard deviation between assays.

### FDA-approved drugs blocked PRL-3 induced cell migration in human cancer cells

PRL-3 is well-established to promote cell migration. Scratch assays were used to test whether broad PRL inhibitors would alter the migratory phenotype of PRL-3 overexpressing HEK293T cells and human colorectal cancer cells that have endogenously high levels of PRL-3. Transfection of a PRL-3 expression vector enhanced cell migration in HEK293T cells by 40% (*p* = 0.02, Supplemental Fig. 6). Empty vector and PRL-3 transfected cells were treated with the broad PRL inhibitors at concentrations that were previously found to not affect cell viability by more than 25% (Fig. 3a). Treatment with two of the drugs, Salirasib and Candesartan, resulted in a significant inhibition of cellular migration in PRL-3 overexpressing cells, compared to DMSO treatment (*p* = 0.013 and p = 0.004, Fig. 4a-b), suggesting that their effects originated from PRL-3 inhibition. Other drugs, including Idasanutlin, YM155, and Napabucasin had less significant effects on the migration of PRL-3 expressing cells, compared to control cells, and were therefore excluded from further analysis (Supplemental Fig. 7). Next, we tested whether Salirasib and Candesartan could decrease cellular migration in colorectal cancer cells that express high levels of endogenous PRL-3^11,21^. We found that both Salirasib and Candesartan significantly inhibited the migration of colorectal cancer cells by >30% (Hct116, p= 0.03; SW480 p = 0.02; Fig. 4c-e), and that this effect was similar to that seen with the specific PRL inhibitor Theinopyridone. Additionally, neither Salirasib nor Candesartan had a negative effect on the activity of 18 other phosphatases (Supplemental Table 2), suggesting that these drugs selectively target the PRL family over other phosphatases. Taken together these data demonstrate that current FDA-approved drugs can inhibit PRL-3 and block one of its main functional effects in cancer cells.

**Figure 4.**
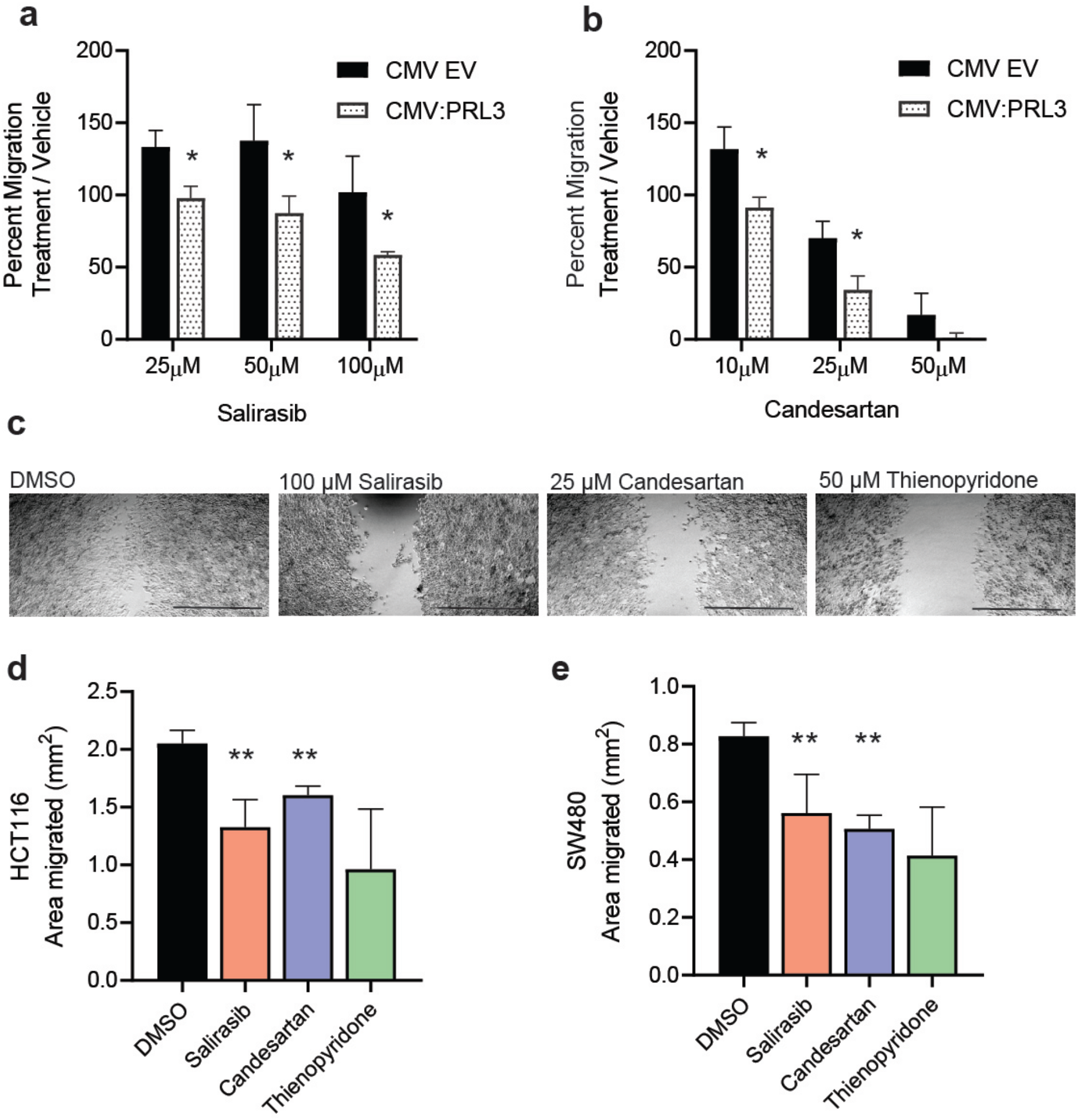
Salirasib and Candesartan inhibit PRL-3 induced cell migration. Quantification of HEK293T cell migration in a scratch assay after treatment with (**a**) Salirasib and (**b**) Candesartan. Both drugs inhibit cell migration to a greater extent in the PRL-3 overexpressing cells, compared to empty vector (EV) control. (**c**) Representative images of migration of the human colorectal cancer cell line Hct116 when treated with DMSO, 100 μM Salirasib, 25 μM Candesartan or 50 μM Thienopyridone. Quantification of (**d**) Hct116 and (**e**) SW480 migration. Migration was measured as the area units migrated 24 hours after scratch. Percent migration was measured as the area migrated 24 hours after scratch normalized to the area migrated by DMSO control. All assays were run in duplicate wells, in 3 independent experiments. Data represent mean phosphatase activity and error bars represent standard deviation between assays. *p < 0.05 comparing migration CMV EV and CMV:PRL-3 by two tailed student t test with Holm Sidak correction. **p < 0.05 comparing migration with drug to DMSO treatment using one way ANOVA with Tukey HSD.

### In silico docking of Salirasib and Candesartan on PRL-3

Due to the lack of currently available structure of monomeric PRLs in complex with any inhibitor, inhibitor binding sites are largely unknown. Attempts by other groups to obtain co-crystals of PRL-3 with inhibitors have been unsuccessful. Thus, molecular docking was employed to identify the potential binding sites of drugs, including the potent inhibitors (Salirasib and Candestartan) in our assays, and to further understand their mechanisms of action. The docking simulations show that, expect for Hexachlorophene, all of the drugs bind preferentially (with lower binding energies) to the closed conformation of PRL-3 (Table 2, Supplemental Fig. 8). In the closed form, two major binding sites were identified: Site 1, which is flanked by the WPD loop and helices α3, α4, and α6 and has been previously predicted to be an allosteric binding site for the research-grade PRL-3 inhibitor, JMS-053^11^, and Site 2, a potential secondary allosteric site adjacent to the PRL-3 CX_5_R motif and bound by helix α4 and the ß-sheets (Fig. 5a). Lineweaver Burke analyses show that both Candestartan and Salirasib reduce apparent V_max_ but do not affect the K_M_ and are non-competitive inhibitors to the substrate (Fig. 5b-e). Therefore, both inhibitors bind sites other than the active site, which is consistent with our *in silico* models indicating that both compounds bind to potential allosteric sites (Supplemental Fig. 9).

**Table 2:**
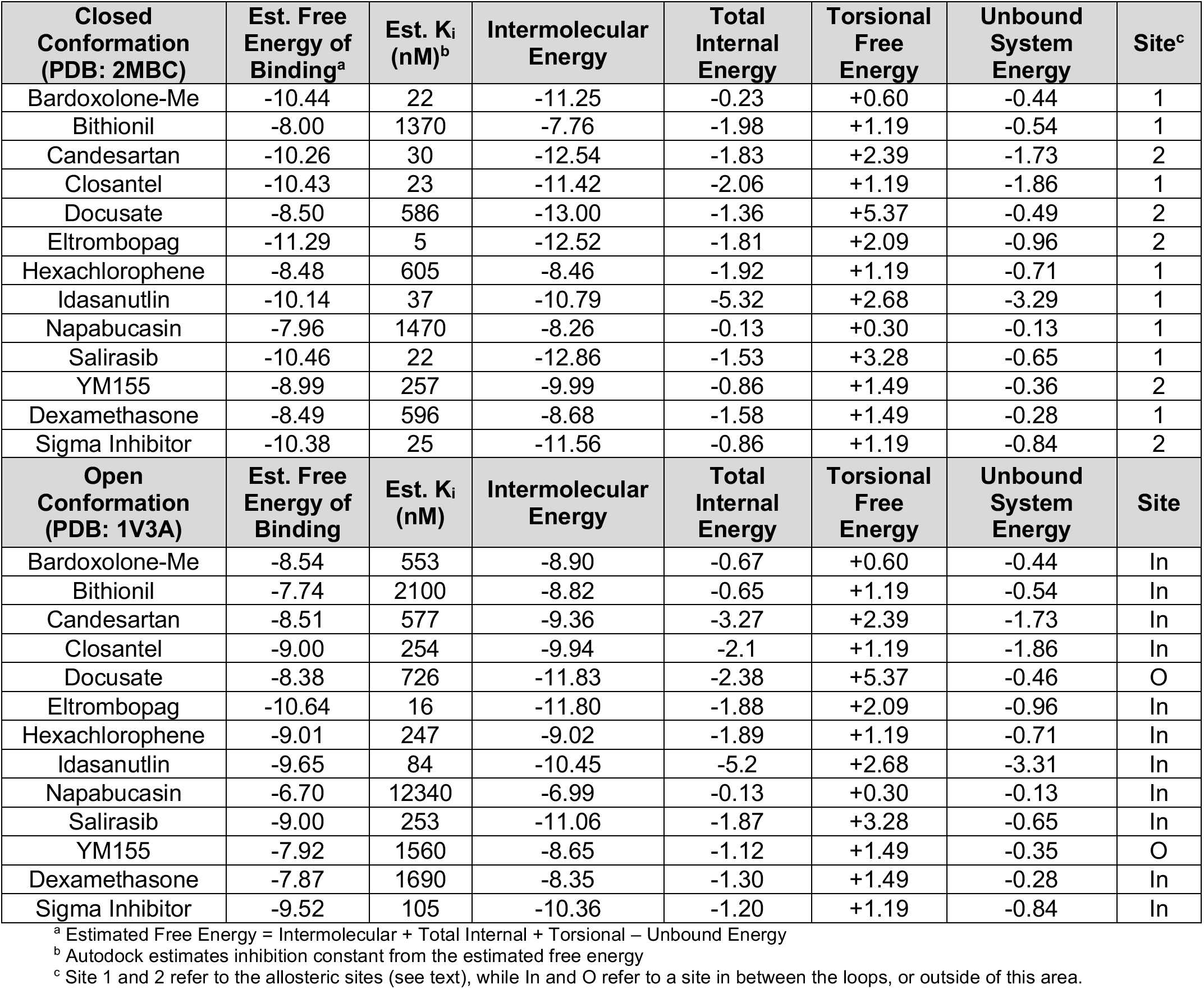
Docking results of FDA-approved drugs with the phosphatase PRL-3 in the open and closed forms.

| **Closed Conformation (PDB: 2MBC)** | **Est. Free Energy of Binding<sup>a</sup>** | **Est. K<sub>i</sub> (nM)<sup>b</sup>** | **Intermolecular Energy** | **Total Internal Energy** | **Torsional Free Energy** | **Unbound System Energy** | **Site<sup>c</sup>** |
| --- | --- | --- | --- | --- | --- | --- | --- |
| Bardoxolone-Me | -10.44 | 22 | -11.25 | -0.23 | +0.60 | -0.44 | 1 |
| Bithionil | -8.00 | 1370 | -7.76 | -1.98 | +1.19 | -0.54 | 1 |
| Candesartan | -10.26 | 30 | -12.54 | -1.83 | +2.39 | -1.73 | 2 |
| Closantel | -10.43 | 23 | -11.42 | -2.06 | +1.19 | -1.86 | 1 |
| Docusate | -8.50 | 586 | -13.00 | -1.36 | +5.37 | -0.49 | 2 |
| Eltrombopag | -11.29 | 5 | -12.52 | -1.81 | +2.09 | -0.96 | 2 |
| Hexachlorophene | -8.48 | 605 | -8.46 | -1.92 | +1.19 | -0.71 | 1 |
| Idasanutlin | -10.14 | 37 | -10.79 | -5.32 | +2.68 | -3.29 | 1 |
| Napabucasin | -7.96 | 1470 | -8.26 | -0.13 | +0.30 | -0.13 | 1 |
| Salirasib | -10.46 | 22 | -12.86 | -1.53 | +3.28 | -0.65 | 1 |
| YM155 | -8.99 | 257 | -9.99 | -0.86 | +1.49 | -0.36 | 2 |
| Dexamethasone | -8.49 | 596 | -8.68 | -1.58 | +1.49 | -0.28 | 1 |
| Sigma Inhibitor | -10.38 | 25 | -11.56 | -0.86 | +1.19 | -0.84 | 2 |
| **Open Conformation (PDB: 1V3A)** | **Est. Free Energy of Binding** | **Est. K<sub>i</sub> (nM)** | **Intermolecular Energy** | **Total Internal Energy** | **Torsional Free Energy** | **Unbound System Energy** | **Site** |
| Bardoxolone-Me | -8.54 | 553 | -8.90 | -0.67 | +0.60 | -0.44 | In |
| Bithionil | -7.74 | 2100 | -8.82 | -0.65 | +1.19 | -0.54 | In |
| Candesartan | -8.51 | 577 | -9.36 | -3.27 | +2.39 | -1.73 | In |
| Closantel | -9.00 | 254 | -9.94 | -2.1 | +1.19 | -1.86 | In |
| Docusate | -8.38 | 726 | -11.83 | -2.38 | +5.37 | -0.46 | O |
| Eltrombopag | -10.64 | 16 | -11.80 | -1.88 | +2.09 | -0.96 | In |
| Hexachlorophene | -9.01 | 247 | -9.02 | -1.89 | +1.19 | -0.71 | In |
| Idasanutlin | -9.65 | 84 | -10.45 | -5.2 | +2.68 | -3.31 | In |
| Napabucasin | -6.70 | 12340 | -6.99 | -0.13 | +0.30 | -0.13 | In |
| Salirasib | -9.00 | 253 | -11.06 | -1.87 | +3.28 | -0.65 | In |
| YM155 | -7.92 | 1560 | -8.65 | -1.12 | +1.49 | -0.35 | O |
| Dexamethasone | -7.87 | 1690 | -8.35 | -1.30 | +1.49 | -0.28 | In |
| Sigma Inhibitor | -9.52 | 105 | -10.36 | -1.20 | +1.19 | -0.84 | In |

<sup>a</sup> Estimated Free Energy = Intermolecular + Total Internal + Torsional – Unbound Energy
<sup>b</sup> Autodock estimates inhibition constant from the estimated free energy
<sup>c</sup> Site 1 and 2 refer to the allosteric sites (see text), while In and O refer to a site in between the loops, or outside of this area.

**Figure 5:**
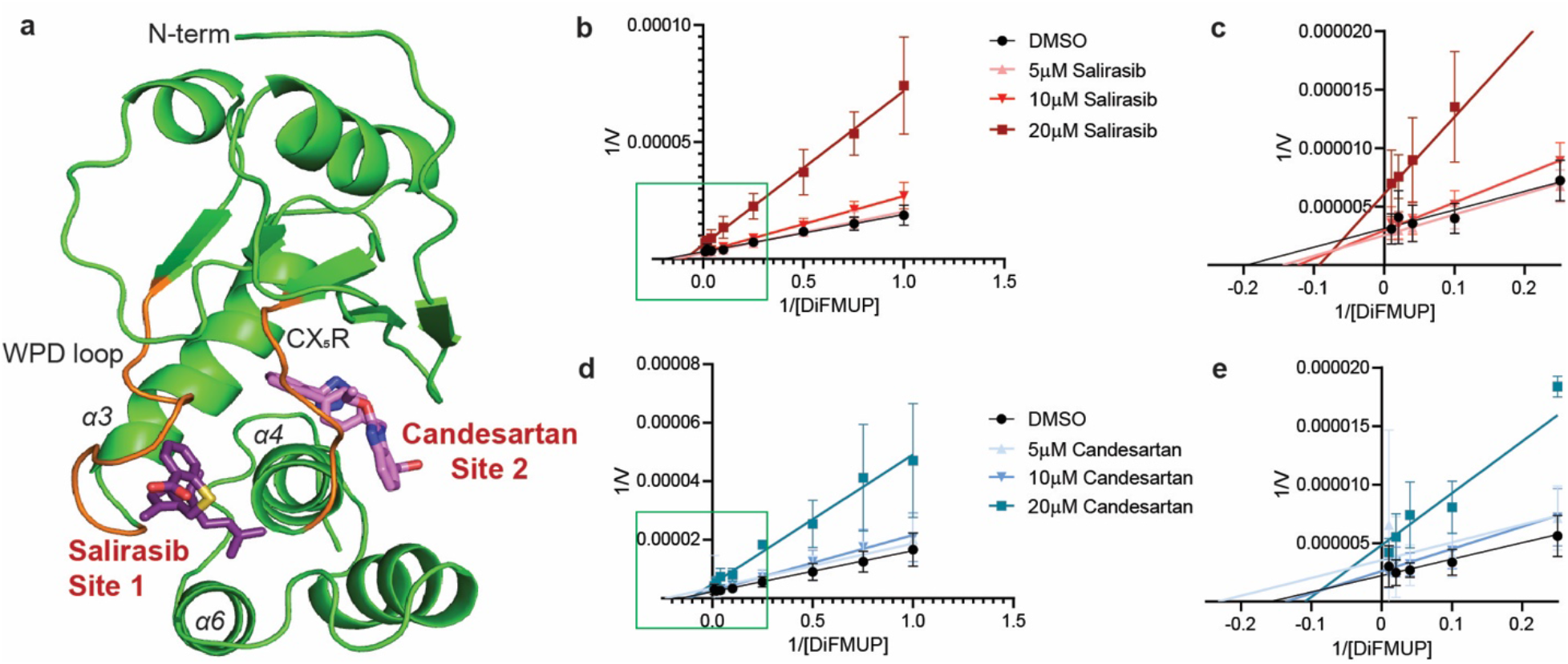
Molecular docking experiments reveal that top hits from screening FDA-approved drugs bind to allosteric sites. (**a**) Representive compounds, Salirasib and Candesartan, are shown bound to two identified allosteric sites adjacent to the active site looops (orange) in the closed conformation of PRL-3. Line-weaver burk plots indicate that Salirasib (**b,c**) and Candesartan (**d**,**e**) are non-competitive inhibitors to the substrate, supporting the docking results. Graphs in **c** and **e** are insets of the boxed areas in **b** and **d**. Assays were run with technical duplicates and performed in 3 independent replicates. Error bars represent standard deviation between assays.

Salirasib binds in the allosteric Site 1 with a free energy of binding of −10.46 kcal/mol. This pocket is lined with hydrophobic residues that likely stabilizes the farnesyl tail of the drug. The salicylic acid group of Salirasib is then free to form hydrogen bonding, in this case with the backbone amide and carbonyl groups of G73 (Fig. 6a,c), likely stabilizing PRL-3 in a closed conformation. Candesartan binds at the potentially new allosteric Site 2 with an estimated free energy of binding of −10.26 kcal/mol for the closed site. In particular, it fits in a shallow binding pocket lined with hydrophobic residues, similar to Site 1, and can form electrostatic interactions with PRL-3 through the sidechain of K136 and the backbone carbonyl of A106 (Fig. 6b,d). The low docking scores are expected due to the low molecular weight of the drugs. The drugs and the sites are also largely hydrophobic. Regardless, these docking simulations show a potential mechanism for PRL-3 inhibition by Salirasib and Candesartan. Additionally, Site 2 offers an additional targetable site that to our knowledge has not been pursued specifically in the past.

**Figure 6.**
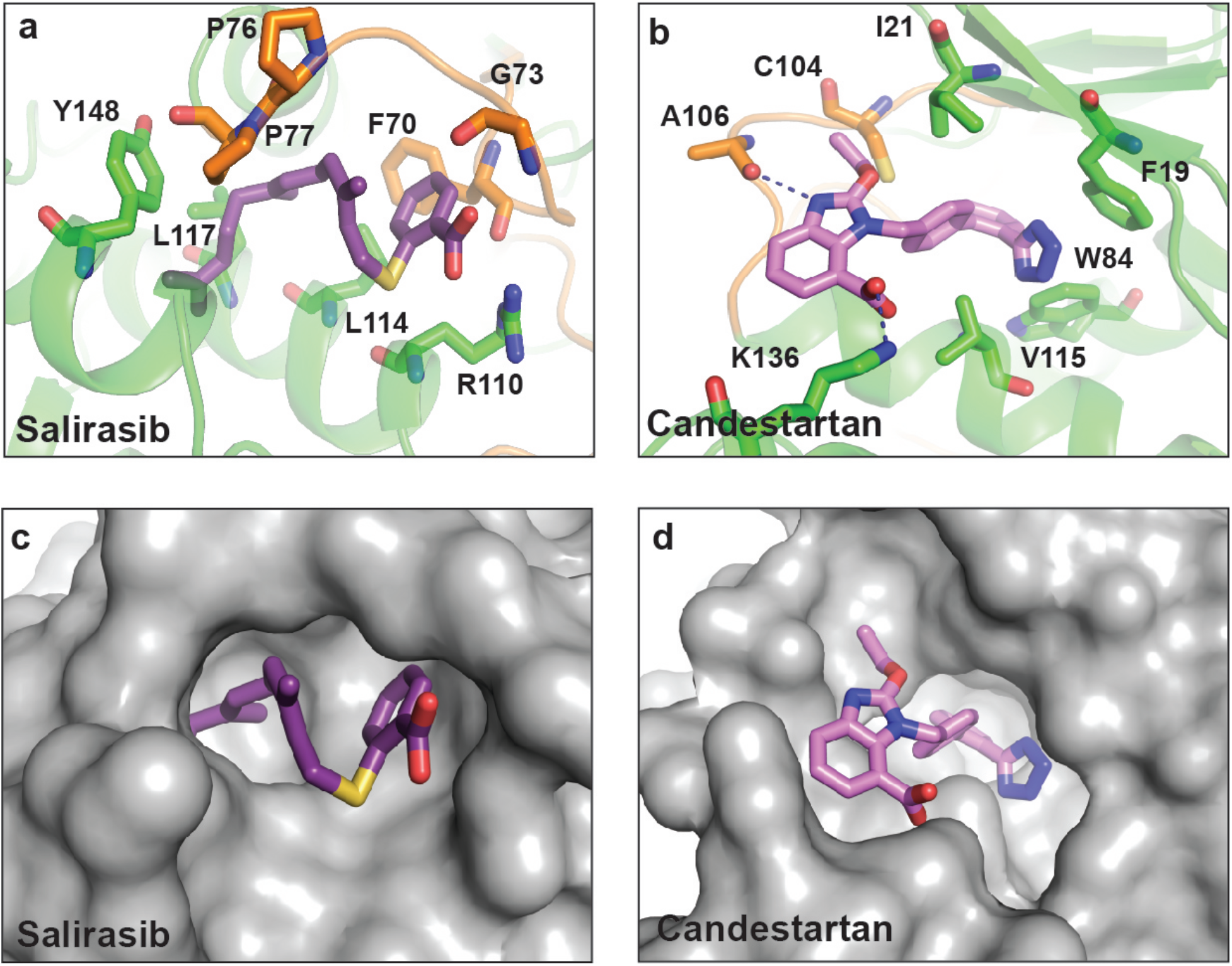
Proposed binding modes for Salirasib and Candesartan with PRL-3. Salirasib (**a**) and Candesartan (**b**) are shown bound to the closed conformation of PRL-3 (PDB: 2MBC) with the active site loops colored in orange. Residues in close proximity to the bound molecule are identified. Surface representation of PRL-3 reveal how Salirasib (**c**) and Candesartan (**d**) fit into pockets within the closed conformation.

As a validation of these results, drugs that were used as controls in experimental work were docked. Dexamethasone was used as a negative control, while the rhodanine derivative PRL-3 Inhibitor (Sigma) was used as a positive control in the drug screen. In both conformations, dexamethasone the lowest ranked, while PRL-3 Inhibitor is among the higher ranked. Finally, a farnesyl tail was docked to verify binding of the farnesyl derivative, Salirasib. Blind docking identified the same binding pocket in Site 1 (Supplemental Fig. 10).

## Discussion

PRL-3 is highly expressed in many cancer types and is a proven oncoprotein. It has well-established roles in tumor cell invasion and metastasis, and data suggests it may also be involved in cancer cell proliferation and drug resistance^47,48^. There have been increasing efforts to identify PRL-3 inhibitors. A screen of the Korea Chemical Bank identified the rhodanine derivatives CG-707 and BR-1 as PRL inhibitors, but further analysis showed these to be fairly nonspecific^43^. Screening of the Roche chemical library identified Thienopyridone as a PRL-3 inhibitor that reduced tumor growth by interfering with cell adhesion, although it is too toxic for *in vivo* use^39^. Further analysis of Thienoypridone, relying on active site mimicry and *in silico* structural modeling, led to the discovery of additional PRL-3 inhibitor Analog 3^40^. A subsequent structural activity relationship study identified iminothienopyridinedione 13 (JMS-053), an analog with lower toxicity and increased potency than Theinopryidone^41^. This compound is notable because it is the first PRL-3 small molecule inhibitor used *in vivo*, where it inhibited cancerous cellular growth in a multi-drug resistant ovarian xenograft model. However, the solubility of JMS-053 is poor, and it requires further study to determine pharmacokinetics, toxicity, and efficacy in humans. Finally, a specific PRL-3 binding antibody has been developed, which is currently the only inhibitor capable of selectively targeting PRL-3 over other family members and is capable of preventing tumor growth using *in vivo* model of both hepatic and gastric cancers^15,49^. Although antibody treatments are an effective strategy for targeting proteins in a variety of diseases, the associated costs are significantly higher than using a small molecule inhibitor, and this antibody will also require clinical trials for safety and efficacy. Although efforts are underway to improve these compounds, it can take many years to bring a drug from development to the clinic.

Currently, there are no available structures of PRL-3 in complex with any inhibitor. This lack of information makes it challenging to define residues in PRL-3 and functional groups in the compounds that are key to the PRL-3-inhibitor interaction for optimization efforts. Without these structures, it is also difficult to identify targetable sites in PRL-3 or propose binding sites for potential inhibitors. Through our screen of >1,400 FDA-approved drugs, we identified several compounds that have potential off-label application as PRL inhibitors, which were predicted to bind at two distinct allosteric sites. Salirasib, also known as farnesylthiosalicylic acid, binds to a previously identified binding pocket adjacent to the WPD loop. The allosteric inhibitor JMS-053 was also proposed to bind to this site^11^. Meanwhile, Candesartan binds to a secondary pocket at the opposite side of the active site, adjacent to the CX_5_R motif. Both of these binding sites are lined with hydrophobic residues that may allow binding of highly hydrophobic small molecules, like Candesartan (logP of 6.1) and Salirasib (logP of 6.8).

The WDP loops and the CX_5_R motif (or P loop) harbor the catalytic residues of classical dual-specificity phosphatases. The two allosteric sites identified are located adjacent to these loops, and drug binding might be sensitive to PRL-3 conformation, as both drugs, as well as most others tested, have lower binding energies with the closed conformation. The catalytic cycle of PRLs and other classical phosphatases are characterized by large structural rearrangements primarily in the WPD loop and surrounding region. As PRL-3 transitions to its open state, the catalytic loops move a total of about 10Å towards each other, moving the catalytic residues into close proximity. This includes the nucleophilic cysteine (P loop, C104 in PRL-3), the aspartic acid (WPD loop, D72) that acts as the general base/proton donor, and the conserved arginine (P loop, R110) that stabilizes the phosphoenzyme intermediate. The preference for the closed conformation indicates a possible mechanism of action where these small molecules trap PRL-3 in the closed conformation, preventing active site structural rearrangements, which is necessary for catalysis^50,51^ Finally, to our knowledge, the secondary allosteric site that was identified for Candesartan and others (Figure 5, Supplemental Table 2) has not been previously identified before as a binding site for inhibitors of PRL-3 and provides an alternative target site for *in silico* screens to identify novel molecules targeting PRL-3. Binding of Salirasib and Candesartan to potential allosteric sites is supported by our enzyme kinetics studies, which show that both are non-competitive inhibitors. Additionally, while the FDA-approved drugs were able to inhibit PRL-3 phosphatase activity and function in cellular migration, the IC_50_s were in the μM range. Future work modifying these compounds may be useful to increase the strength of binding and improve their potency.

Targeting protein phosphatases such as PRL family members has been challenging due to high sequence similarity and the conserved active sites among protein tyrosine phosphatases. While atomic resolution details on how PRLs engage their substrates are not yet available partly due to lack of verified substrates, all PRLs have been crystalized bound to CNNM CBS-pair domains, which show that PRLs engage this pseudo-substrate very similarly^52^. Additionally, small molecule inhibitors have little selectivity since the PRL active site is wide and shallow; the most effective PRL-3 inhibitors identified in this study also targeted the entire PRL family. Allosteric inhibition of PLRs provides an alternate strategy for the identification and development of more specific PRL-3 inhibitors. Thus far, two potential allosteric sites are identified for PRL-3 and further studies are needed to experimentally validate both sites, as well as the possibility of targeting them with small molecules. Several drugs, including Bardoxolone and Eltromobag, showed greater specificity for PRL-3 over PRL-1 and PRL-2. The properties of these drugs may be useful in informing future development of novel PRL-3 inhibitors.

## Methods

### Recombinant protein expression

PRL full length protein sequences fused to a C-terminal 6xHis-tag was cloned into pET28b bacterial expression vector. Proteins were expressed in One Shot^TM^ BL21 Star^TM^ bacteria (Invitrogen Cat No. C601003) by induction with 0.5 mM IPTG (Fisher Scientific Cat. No. BP175510) for 16hours. Cells were resuspended 10ml of lysis buffer [300 mM NaCl (VWR Cat. No. BDH9286), 20mM Tris pH 7.5, 10 mM Imidazole pH 8.0 (Sigma-Aldrich I2399), 1:1000 protease inhibitor cocktail (Sigma-Aldrich Cat. No. P8465)] per gram of cell pellet and lysed using a microfluidizer (Avestin, EmulsiFlex-C5). Protein was isolated using Ni-NTA Resin (VWR Cat. No. 786-940) and eluted with 2 mL of elution buffer (300 mM NaCl, 20 mM Tris pH 7.5, and 250 mM Imidazole pH 8.0). Following cleavage with TEV protease, samples were reapplied to Ni-NTA column to remove uncleaved protein as well as TEV. Samples were further purified using an S200 column on a Superdex 10/300 in buffer containing 100mM NaCl and 200 mM HEPES pH 7.5. Purified fractions were then run on 4-20% Mini-PROTEAN TGX Stain-Free^TM^. The purest fractions were pooled, concentrated together, flash frozen and stored at −80°C.

### Drug Panel and Other Reagents

The library of FDA-approved drugs (Selleck, L1300) was generously provided by Dr. Vivek Rangnekar at the University of Kentucky. For further testing, additional drugs were purchased as listed in Supplemental Table 3. Thienopyridone was generously provided by Dr. Zhong-Yin Zhang (Purdue University). PRL-3 over expressing construct was made by cloning full length PRL-3 cDNA into p3XFLAG-CMV-14 expression vector (Sigma, E7908)

### *In vitro* phosphatase assay

In 384 well plates, 2.5 μM recombinant PRL-1, PRL-2, or PRL-3 was combined with DiFUMP (Life Technologies, E12020) at the K_M_ of each protein in reaction buffer (20 mM Tris-Cl, pH 7.5, 150 mM NaCl, 10 mM DTT) as previously reported^40^. Briefly, protein was diluted in reaction buffer and allowed to incubate at 4° C for 20 minutes to allow for full reduction of the active site. DiFMUP, drug, and protein were added, and plates were incubated for 20 minutes at room temperature. Fluorescence intensities were measured on a Cytation 5 plate reader (Biotek) at an EX:360 nm and EM:460 nm. Data were normalized to vehicle control and IC50 values were calculated on GraphPad Prism version 8. Inhibition selectivity for PRLs over a panel of other phosphatases was determined using the PhosphataseProfilerTM (Eurofins, PP260).

### Cell culture

HEK293T, Hct116, and SW480 human cell lines, from ATCC, CRL-3216, CCL-247, CCL-228, respectively, were maintained in 1X DMEM with glutamine and glucose (Gibco, 11965-092) supplemented with 10% FBS (Atlanta Biologicals, S11150H) and Penicillin/Streptomycin (Fisher Scientific, 10378016). Cells were cultured at 37°C in a humidified incubator with 5% CO_2_. Transfection was preformed using Lipofectamine 3000 (ThermoFisher, L3000015) following manufacturer’s protocol.

### Scratch Assay

Cells were seeded into duplicate wells of a 48 well plate at a density of 2×10^5^ cells per well for HEK293T or 4×10^5^ for Hct116 and SW480. Cells recovered overnight, and were scratched using a p20 pipette tip the following day. Compounds were added at the concentrations indicated. Images were obtained immediately after the scratch and after a 24 hour incubation with the drug on an Evos FL inverted microscope (ThermoFisher). The area within the scratch was calculated using ImageJ. The area migrated was measured as the difference in wound area at 0h and 24h, and percent migration represents area migrated in the drug treated cells divided by the area migrated in vehicle control.

### Viability Assay

Cells were seeded into duplicate wells of a 96 well plate at a density of 2×10^4^ cells per well and allowed to recover for 24 hours. Cells were then treated with drug or DMSO at the indicated concentrations for 16 hours. Following drug treatment, 10 μL of 5 mg/ml MTT (Sigma, M5655) was added to each well and cells were incubated for 4 hours. Finally, media were removed, and the dye was solubilized in 100 μL of 0.1M HCl in isopropanol. Absorbance was measured at 570nm and 690nm with final absorbance measurements as 570nm-690nm.

### Western Blot

Cells were lysed using Qproteome lysis buffer, then spun at 14000K RPM for 10 minutes at 4°C to pellet debris (Qiagen, 37901). Protein concentration in the supernatant was quantified using BCA assay (Thermo Scientific, 23227). 30 μg of protein was loaded into each lane of a TGX-stain-free pre-cast 4–20% SDS gel (Biorad, 4568094), total protein was quantified upon stain-free gel imaging, and protein was transferred onto PVDF membrane using the Trans-Blot Turbo Transfer System (BioRad, 1704150). Membranes were blocked with 5% milk in 1% TBST for 1 hour, and a 1:1000 dilution of Anti PRL-3 antibody (R&D Biosystems, MAB3219) was added overnight. Following three washes in TBST, secondary HRP-conjugated antibody (GE, NA9340V) were added at a 1:5000 dilution for 1 hour and membranes imaged using Clarity Western ECL Substrate. (BioRad, 1705061).

### Molecular docking

The molecular docking software, Autodock41-3 (version 4.2.6) was used to identify probable binding sites. Protein structures were obtained from published NMR solution structures of apo (PDB code: 1V3A4) and vanadate-bound (PDB code: 2MBC5, model 1) PRL-3. Ligand sdf files (pubchem.com6) were converted to PDB using OpenBabel7.8. Ligand and receptor files were prepared and Gasteiger charges added using AutodockTools1. Blind Autodock docking was performed by covering the entire protein structure with a grid box consisting of 126 × 126 × 126 points and centered at the center of the macromolecule. Docking was then performed without bias to search for all possible binding sites using Lamarckian Genetic Algorithm with a rigid protein and flexible ligands. Population size was set to 300 and the maximum number of energy evaluations and maximum number of generations were set to 30,000,000 and 27,000, respectively. For each ligand, 100 dockings were performed and clustered using an RMSD tolerance of 2Å. In addition to the set of FDA-approved drugs, dexamethasone and PRL inhibitor I, which were used as negative and positive controls in experimental work, respectively, as well as a farnesyl tail were docked following the same protocols. Visualizations were performed using AutodockTools and PyMol10.

### Statistics

All experiments, expect for the drug screen, were performed in biological triplicate across at least three independent time points with at least two technical replicates. Where applicable, experimental values were normalized to vehicle control, ± standard deviation. Unless otherwise noted p values were calculated using one way ANOVA with Tukey HSD in Graphpad Prism Version 8, and changes were considered significant if p < 0.05.

## Supporting information

All Supplemental Figures

## Data Availability

All data generated or analyzed during this study are included in this article and its supplementary files. The datasets that were analyzed are available from the corresponding authors on reasonable request.

## Acknowledgements

We thank Vivek Ragnakar for providing the FDA approved library, and Zhong-Ying Zhang for providing Thienopryridone. We thank Konstantin Korotkov for assistance with generating Pymol structures. This research was supported by a St. Baldrick’s Foundation Research Grant, and NIH grants DP2CA228043 and R37CA227656 (to J.S. Blackburn). The research was also supported by the James Graham Brown Cancer Center (to D. Lee)

## Author Contributions

D.R.R. and J.S.B. conceived of and designed the study. D.R.R. performed all biochemical and cell-based experiments, with B.P. assisting on some assays. M.D.C and D.L. designed and carried out molecular docking analyses and wrote the corresponding portion of the manuscript. C.N.S. generated all protein used in this study. D.R.R. and M.D.C. drafted the manuscript, J.S.B and D.L. revised. All authors have read, edited, and approved of the final version of this manuscript.

## Competing Interests

The authors declare not conflicts of interest.

## Notes

### Competing Interest Statement

The authors have declared no competing interest.

## References

1 Ruckert, M. T., de Andrade, P. V., Santos, V. S. & Silveira, V. S. Protein tyrosine phosphatases: promising targets in pancreatic ductal adenocarcinoma. Cell Mol Life Sci 76, 2571–2592, doi:10.1007/s00018-019-03095-4 (2019).

2 Aceto, N. et al. Tyrosine phosphatase SHP2 promotes breast cancer progression and maintains tumor-initiating cells via activation of key transcription factors and a positive feedback signaling loop. Nat Med 18, 529–537, doi:10.1038/nm.2645 (2012).

3 Hoekstra, E. et al. Increased PTP1B expression and phosphatase activity in colorectal cancer results in a more invasive phenotype and worse patient outcome. Oncotarget 7, 21922–21938, doi:10.18632/oncotarget.7829 (2016).

4 Hu, Z., Li, J., Gao, Q., Wei, S. & Yang, B. SHP2 overexpression enhances the invasion and metastasis of ovarian cancer in vitro and in vivo. Onco Targets Ther 10, 3881–3891, doi:10.2147/ott.s138833 (2017).

5 Lessard, L. et al. PTP1B is an androgen receptor-regulated phosphatase that promotes the progression of prostate cancer. Cancer Res 72, 1529–1537, doi:10.1158/0008-5472.can-11-2602 (2012).

6 Bardelli, A. et al. PRL-3 expression in metastatic cancers. Clin Cancer Res 9, 5607–5615 (2003).

7 Campbell, A. M. & Zhang, Z. Y. Phosphatase of regenerating liver: a novel target for cancer therapy. Expert Opin Ther Targets 18, 555–569, doi:10.1517/14728222.2014.892926 (2014).

8 Dai, N., Lu, A. P., Shou, C. C. & Li, J. Y. Expression of phosphatase regenerating liver 3 is an independent prognostic indicator for gastric cancer. World J Gastroenterol 15, 1499–1505, doi:10.3748/wjg.15.1499 (2009).

9 den Hollander, P. et al. Phosphatase PTP4A3 Promotes Triple-Negative Breast Cancer Growth and Predicts Poor Patient Survival. Cancer Res 76, 1942–1953, doi:10.1158/0008-5472.can-14-0673 (2016).

10 Park, J. E. et al. Oncogenic roles of PRL-3 in FLT3-ITD induced acute myeloid leukaemia. EMBO Mol Med 5, 1351–1366, doi:10.1002/emmm.201202183 (2013).

11 Saha, S. et al. A phosphatase associated with metastasis of colorectal cancer. Science 294, 1343–1346, doi:10.1126/science.1065817 (2001).

12 Vandsemb, E. N. et al. Phosphatase of regenerating liver 3 (PRL-3) is overexpressed in human prostate cancer tissue and promotes growth and migration. J Transl Med 14, 71, doi:10.1186/s12967-016-0830-z (2016).

13 Wang, L. et al. PTP4A3 is a target for inhibition of cell proliferatin, migration and invasion through Akt/mTOR signaling pathway in glioblastoma under the regulation of miR-137. Brain Res 1646, 441–450, doi:10.1016/j.brainres.2016.06.026 (2016).

14 Yeh, H. C. et al. PTP4A3 Independently Predicts Metastasis and Survival in Upper Tract Urothelial Carcinoma Treated with Radical Nephroureterectomy. J Urol 194, 1449–1455, doi:10.1016/j.juro.2015.05.101 (2015).

15 Thura, M. et al. PRL3-zumab as an immunotherapy to inhibit tumors expressing PRL3 oncoprotein. Nat Commun 10, 2484, doi:10.1038/s41467-019-10127-x (2019).

16 Beekman, R. et al. Retroviral integration mutagenesis in mice and comparative analysis in human AML identify reduced PTP4A3 expression as a prognostic indicator. PLoS One 6, e26537, doi:10.1371/journal.pone.0026537 (2011).

17 Mayinuer, A. et al. Upregulation of protein tyrosine phosphatase type IVA member 3 (PTP4A3/PRL-3) is associated with tumor differentiation and a poor prognosis in human hepatocellular carcinoma. Ann Surg Oncol 20, 305–317, doi:10.1245/s10434-012-2395-2 (2013).

18 Qu, S. et al. Independent oncogenic and therapeutic significance of phosphatase PRL-3 in FLT3-ITD-negative acute myeloid leukemia. Cancer 120, 2130–2141, doi:10.1002/cncr.28668 (2014).

19 Ren, T. et al. Prognostic significance of phosphatase of regenerating liver-3 expression in ovarian cancer. Pathol Oncol Res 15, 555–560, doi:10.1007/s12253-009-9153-1 (2009).

20 Guo, K. et al. Catalytic domain of PRL-3 plays an essential role in tumor metastasis: formation of PRL-3 tumors inside the blood vessels. Cancer Biol Ther 3, 945–951, doi:10.4161/cbt.3.10.1111 (2004).

21 Kato, H. et al. High expression of PRL-3 promotes cancer cell motility and liver metastasis in human colorectal cancer: a predictive molecular marker of metachronous liver and lung metastases. Clin Cancer Res 10, 7318–7328, doi:10.1158/1078-0432.ccr-04-0485 (2004).

22 Wu, X. et al. Phosphatase of regenerating liver-3 promotes motility and metastasis of mouse melanoma cells. Am J Pathol 164, 2039–2054, doi:10.1016/s0002-9440(10)63763-7 (2004).

23 Zeng, Q. et al. PRL-3 and PRL-1 promote cell migration, invasion, and metastasis. Cancer Res 63, 2716–2722 (2003).

24 Li, Z. et al. Inhibition of PRL-3 gene expression in gastric cancer cell line SGC7901 via microRNA suppressed reduces peritoneal metastasis. Biochem Biophys Res Commun 348, 229–237, doi:10.1016/j.bbrc.2006.07.043 (2006).

25 Polato, F. et al. PRL-3 phosphatase is implicated in ovarian cancer growth. Clin Cancer Res 11, 6835–6839, doi:10.1158/1078-0432.ccr-04-2357 (2005).

26 Qian, F. et al. PRL-3 siRNA inhibits the metastasis of B16-BL6 mouse melanoma cells in vitro and in vivo. Mol Med 13, 151–159, doi:10.2119/2006-00076.Qian (2007).

27 Zimmerman, M. W., Homanics, G. E. & Lazo, J. S. Targeted deletion of the metastasis-associated phosphatase Ptp4a3 (PRL-3) suppresses murine colon cancer. PLoS One 8, e58300, doi:10.1371/journal.pone.0058300 (2013).

28 Hardy, S. et al. The protein tyrosine phosphatase PRL-2 interacts with the magnesium transporter CNNM3 to promote oncogenesis. Oncogene 34, 986–995, doi:10.1038/onc.2014.33 (2015).

29 Jin, S. et al. Oncogenic function and prognostic significance of protein tyrosine phosphatase PRL-1 in hepatocellular carcinoma. Oncotarget 5, 3685–3696, doi:10.18632/oncotarget.1986 (2014).

30 Liu, L. Z. et al. Protein tyrosine phosphatase PTP4A1 promotes proliferation and epithelial-mesenchymal transition in intrahepatic cholangiocarcinoma via the PI3K/AKT pathway. Oncotarget 7, 75210–75220, doi:10.18632/oncotarget.12116 (2016).

31 Shinmei, S. et al. Identification of PRL1 as a novel diagnostic and therapeutic target for castration-resistant prostate cancer by the Escherichia coli ampicillin secretion trap (CAST) method. Urol Oncol 32, 769–778, doi:10.1016/j.urolonc.2014.03.007 (2014).

32 Wang, Y. & Lazo, J. S. Metastasis-associated phosphatase PRL-2 regulates tumor cell migration and invasion. Oncogene 31, 818–827, doi:10.1038/onc.2011.281 (2012).

33 Dong, J., Sui, L., Wang, Q., Chen, M. & Sun, H. MicroRNA-26a inhibits cell proliferation and invasion of cervical cancer cells by targeting protein tyrosine phosphatase type IVA 1. Mol Med Rep 10, 1426–1432, doi:10.3892/mmr.2014.2335 (2014).

34 Dumaual, C. M. et al. Tissue-specific alterations of PRL-1 and PRL-2 expression in cancer. Am J Transl Res 4, 83–101 (2012).

35 Hardy, S., Wong, N. N., Muller, W. J., Park, M. & Tremblay, M. L. Overexpression of the protein tyrosine phosphatase PRL-2 correlates with breast tumor formation and progression. Cancer Res 70, 8959–8967, doi:10.1158/0008-5472.can-10-2041 (2010).

36 Rios, P., Li, X. & Kohn, M. Molecular mechanisms of the PRL phosphatases. Febs j 280, 505–524, doi:10.1111/j.1742-4658.2012.08565.x (2013).

37 Wei, M., Korotkov, K. V. & Blackburn, J. S. Targeting phosphatases of regenerating liver (PRLs) in cancer. Pharmacol Ther 190, 128–138, doi:10.1016/j.pharmthera.2018.05.014 (2018).

38 Bai, Y. et al. Novel Anticancer Agents Based on Targeting the Trimer Interface of the PRL Phosphatase. Cancer Res 76, 4805–4815, doi:10.1158/0008-5472.can-15-2323 (2016).

39 Daouti, S. et al. A selective phosphatase of regenerating liver phosphatase inhibitor suppresses tumor cell anchorage-independent growth by a novel mechanism involving p130Cas cleavage. Cancer Res 68, 1162–1169, doi:10.1158/0008-5472.can-07-2349 (2008).

40 Hoeger, B., Diether, M., Ballester, P. J. & Kohn, M. Biochemical evaluation of virtual screening methods reveals a cell-active inhibitor of the cancer-promoting phosphatases of regenerating liver. Eur J Med Chem 88, 89–100, doi:10.1016/j.ejmech.2014.08.060 (2014).

41 McQueeney, K. E. et al. Targeting ovarian cancer and endothelium with an allosteric PTP4A3 phosphatase inhibitor. Oncotarget 9, 8223–8240, doi:10.18632/oncotarget.23787 (2018).

42 Salamoun, J. M. & Wipf, P. Allosteric Modulation of Phosphatase Activity May Redefine Therapeutic Value. J Med Chem 59, 7771–7772, doi:10.1021/acs.jmedchem.6b01210 (2016).

43 Ahn, J. H. et al. Synthesis and biological evaluation of rhodanine derivatives as PRL-3 inhibitors. Bioorg Med Chem Lett 16, 2996–2999, doi:10.1016/j.bmcl.2006.02.060 (2006).

44 Souers, A. J. et al. ABT-199, a potent and selective BCL-2 inhibitor, achieves antitumor activity while sparing platelets. Nat Med 19, 202–208, doi:10.1038/nm.3048 (2013).

45 Ding, Q. et al. Discovery of RG7388, a potent and selective p53-MDM2 inhibitor in clinical development. J Med Chem 56, 5979–5983, doi:10.1021/jm400487c (2013).

46 Nakahara, T. et al. YM155, a novel small-molecule survivin suppressant, induces regression of established human hormone-refractory prostate tumor xenografts. Cancer Res 67, 8014–8021, doi:10.1158/0008-5472.can-07-1343 (2007).

47 Hjort, M. A. et al. Phosphatase of regenerating liver-3 is expressed in acute lymphoblastic leukemia and mediates leukemic cell adhesion, migration and drug resistance. Oncotarget 9, 3549–3561, doi:10.18632/oncotarget.23186 (2018).

48 Zhou, J. et al. The pro-metastasis tyrosine phosphatase, PRL-3 (PTP4A3), is a novel mediator of oncogenic function of BCR-ABL in human chronic myeloid leukemia. Mol Cancer 11, 72, doi:10.1186/1476-4598-11-72 (2012).

49 Thura, M. et al. PRL3-zumab, a first-in-class humanized antibody for cancer therapy. JCI Insight 1, e87607, doi:10.1172/jci.insight.87607 (2016).

50 Koveal, D., Clarkson, M. W., Wood, T. K., Page, R. & Peti, W. Ligand binding reduces conformational flexibility in the active site of tyrosine phosphatase related to biofilm formation A (TpbA) from Pseudomonasaeruginosa. J Mol Biol 425, 2219–2231, doi:10.1016/j.jmb.2013.03.023 (2013).

51 Whittier, S. K., Hengge, A. C. & Loria, J. P. Conformational motions regulate phosphoryl transfer in related protein tyrosine phosphatases. Science 341, 899–903, doi:10.1126/science.1241735 (2013).

52 Zhang, H. et al. PRL3 phosphatase active site is required for binding the putative magnesium transporter CNNM3. Sci Rep 7, 48, doi:10.1038/s41598-017-00147-2 (2017).

