## Supplementary material for "A screen of FDA-approved drugs identifies inhibitors of Protein Tyrosine Phosphatase 4A3 (PTP4A3 or PRL-3)": All Supplemental Figures

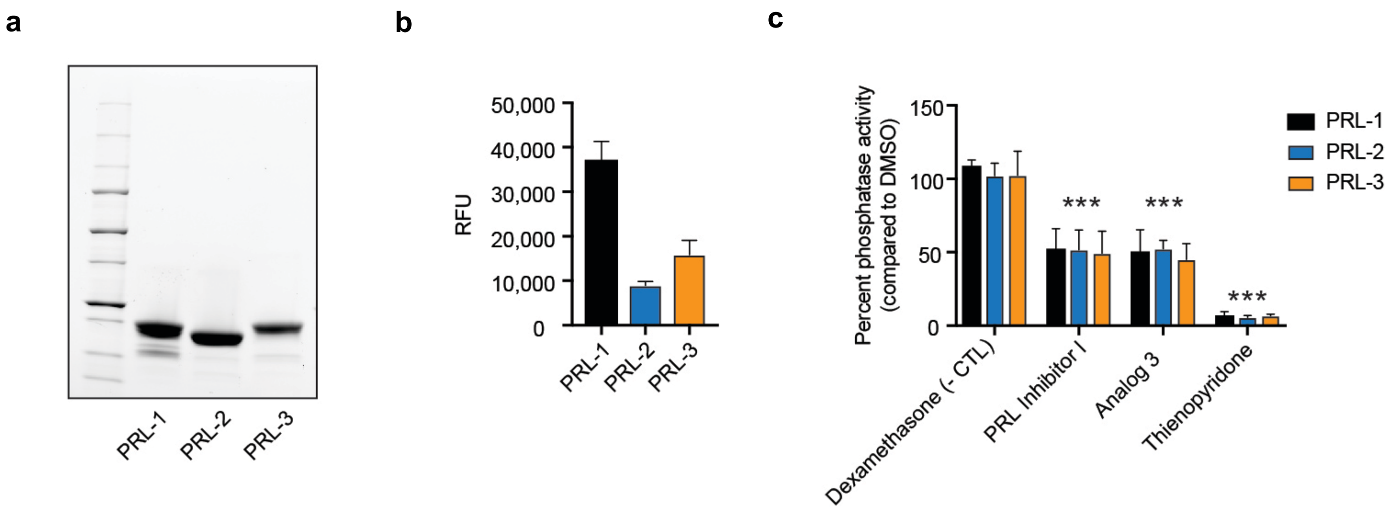


**Supplemental Figure 1. Establishing the efficacy of the DiFMUP assay in assessing PRL phosphatase activity.** A) SDS-PAGE gel demonstrating purity of PRL proteins used in the DiFMUP assay. B) PRL dephosphorylation of DiFMUP measured at ex=360nm and em=460nm. RFU=relative fluorescence units. C) Inhibition of PRL phosphatase activity by known inhibitors, ***p<.0001 compared to the negative control dexamethasone, an FDA-approved drug that is not expected to inhibit PRL activity. Bars represent the average of n=3, and are representative of 3 experiments. Standard deviation is shown.

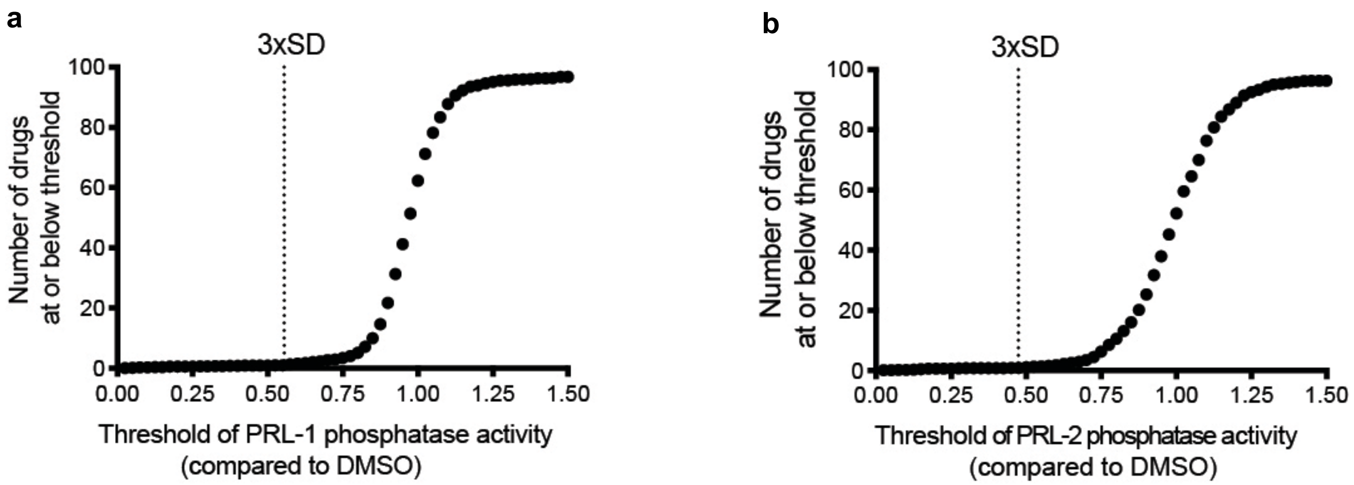


**Supplemental Figure 2. PRL-1 and PRL-2 S-curves.** S-curve representing the positive hit selection strategy of drug screen for PRL-1 (A) and PRL-2 (B). Three-fold standard deviations below the mean phosphatase activity across all drugs is shown (3xSD). Drugs falling to the left of this line were considered hits in this screen.

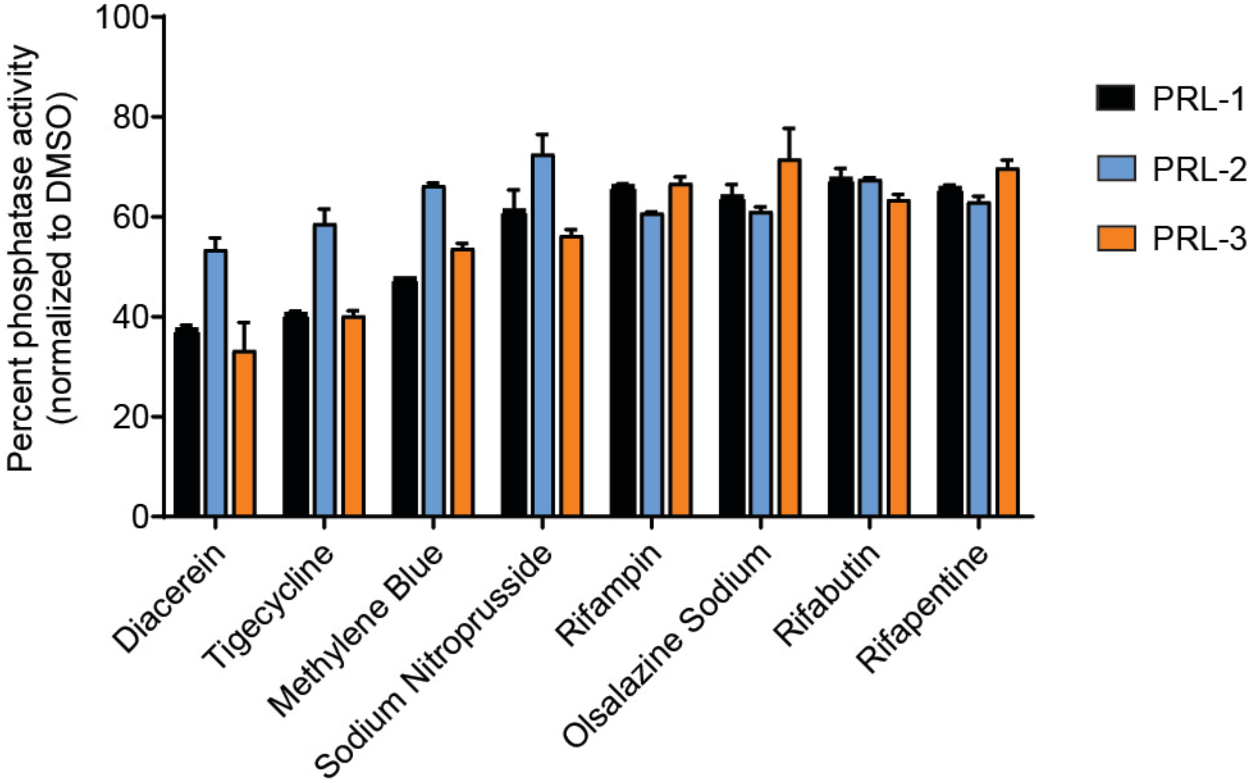


**Supplemental Figure 3. A subset of hits from the initial screen failed re-analysis.** The effects of broad PRL inhibitors on PRL phosphatase activity were examined. While these drugs hit in the initial screen, they did not meet the cut-off of >80% inhibition. Drugs were used at 40μM, bars are average phosphatase activity, the standard deviation is shown.

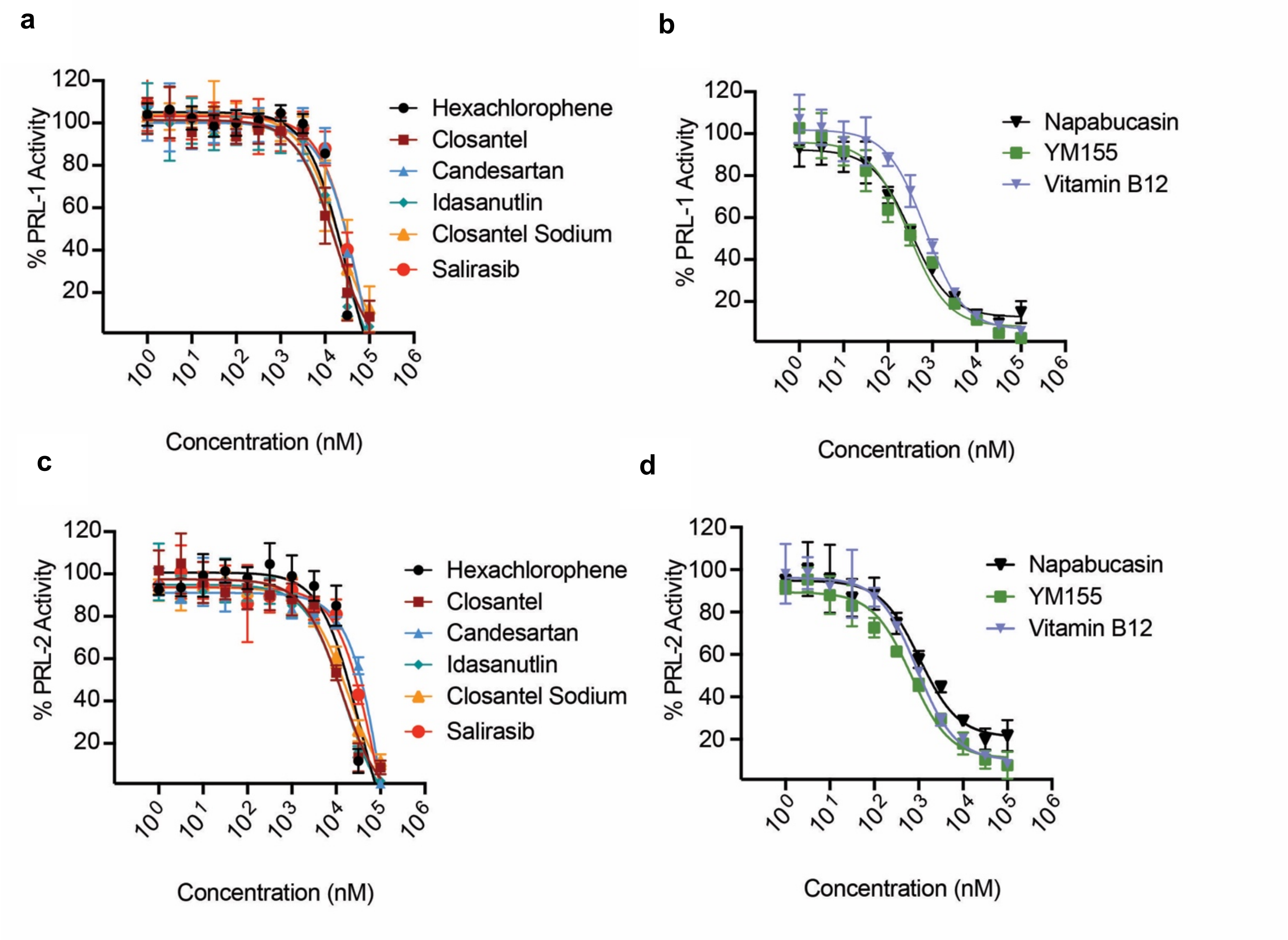


**Supplemental Figure 4. Dose response curves of FDA-approved drugs against PRL-1 and PRL-2.** Dose response curves showing DiFMUP phosphatase activity of PRL-1 (**a,b**) and PRL-2 (**c,d**) when treated with doses ranging between 1 nM and 100 µM of drug. Inhibition of PRL phosphatase activity by the broad PRL family inhibitors are shown, and drugs are subdivided into those with (**a,c**) high $IC_{50}$ values and (**b,d**) low $IC_{50}$ values

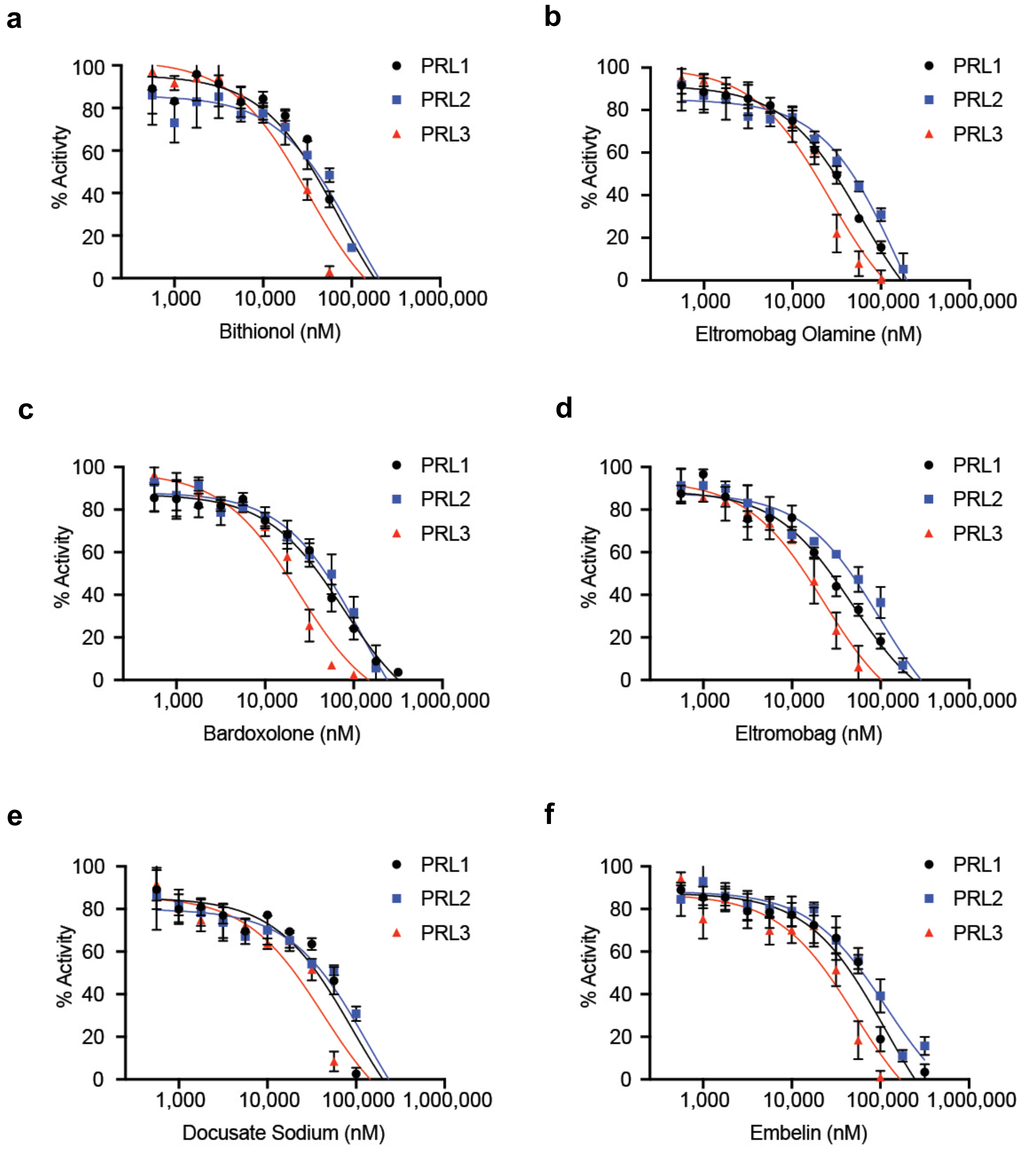


**Supplemental Figure 5. Dose response curves of PRL-3 specific drugs.** (a-f) Inhibition of phosphatase activity by PRL-3 specific compounds. Each data point is the average DiFMUP phosphatase activity of each PRL when treated with the drug dose noted. Assay were run in technical duplicates and are representative of three independent experiments. Standard deviation is shown.

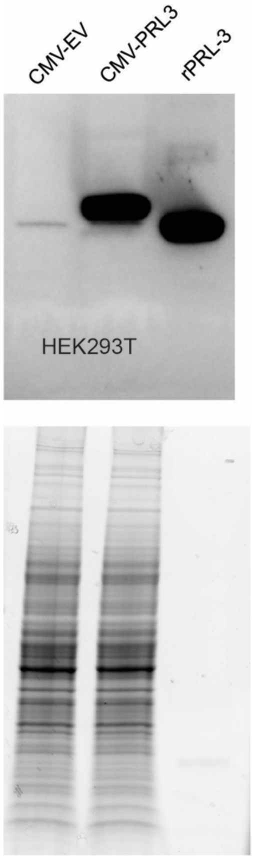


**Supplemental Figure 6. PRL-3 expression in HEK293T.** PRL-3 expression in HEK293T cells after transfection with empty vector control (CMV-EV) or FLAG-PRL-3 overexpressing plasmid (CMV-PRL3). (rPRL-3) was used as a size control, and total protein loaded in the gel is shown.

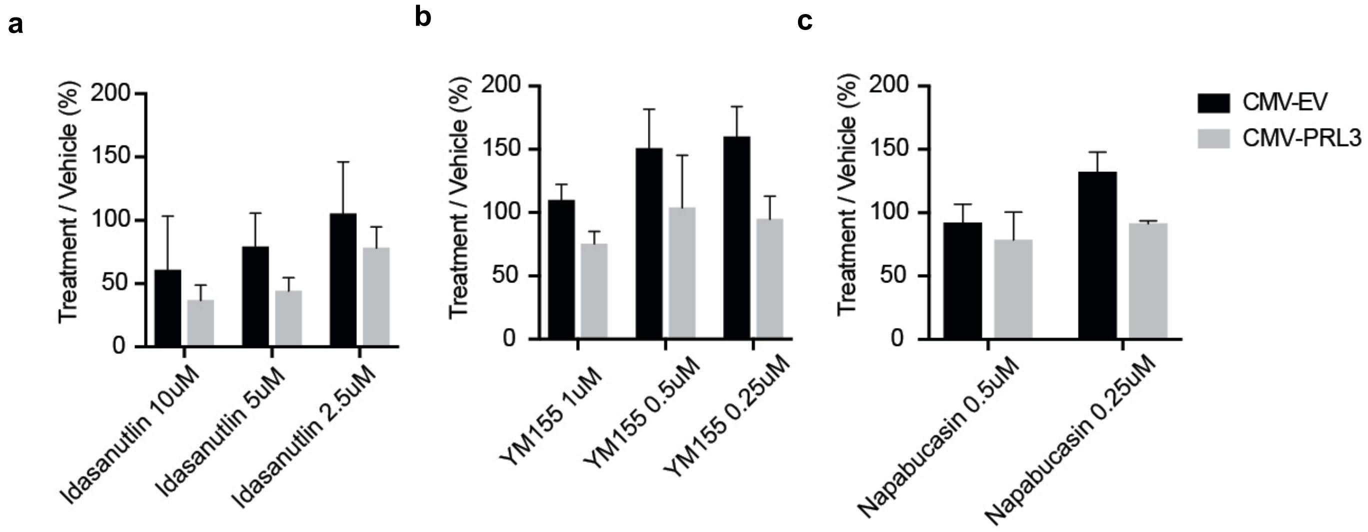


**Supplemental Figure 7. Wound healing assay negative hits.** Migration of vector control and PRL-3 overexpressing HEK293T cells after treatment with (A) Idasanutlin shows the drug impacts migration even in control (EV) cells. Treatment of HEK293T with (B) YM155 and (C) Napabucasin show these drugs do not significantly reduce PRL-3 mediated migration. Migration was measured as the area units migrated 24 hours after scratch. Percent migration was measured as the area units migrated 24 hours after scratch normalized to the area units migrated by DMSO control. All assays were run in duplicate wells, in 3 independent experiments. Statistics were done using one way ANOVA with Tukey HSD.

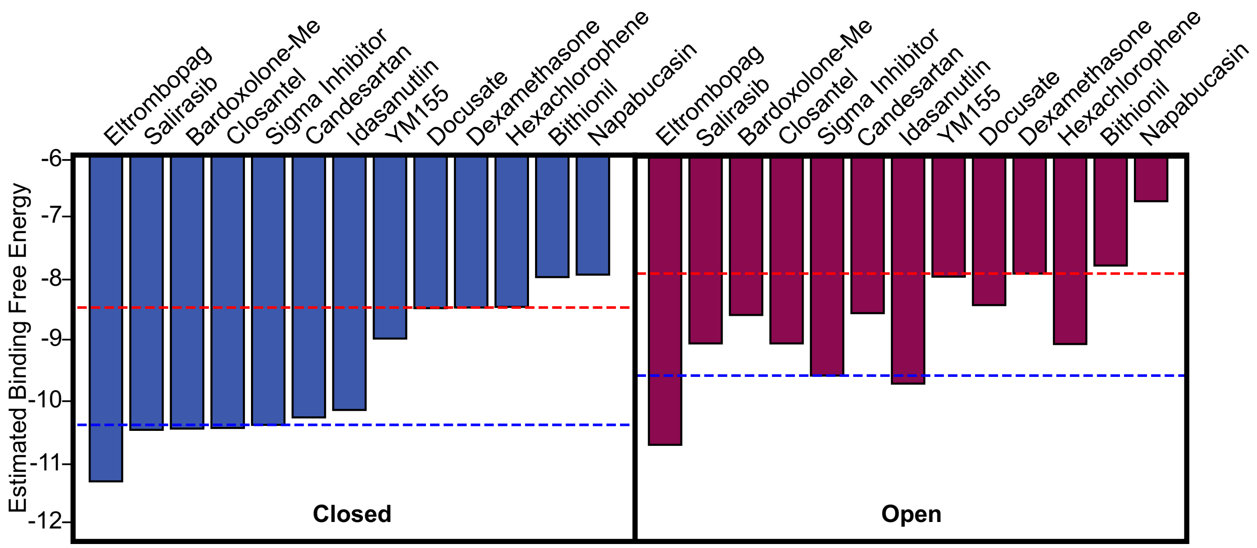


**Supplemental Figure 8. Comparison of estimated free energy of binding of compounds to the open and closed conformation of PRL-3.** Controls are highlighted with Dexamethasone (red dotted line) as negative control and PRL3 Sigma Inhibitor (blue dotted line) as positive control.

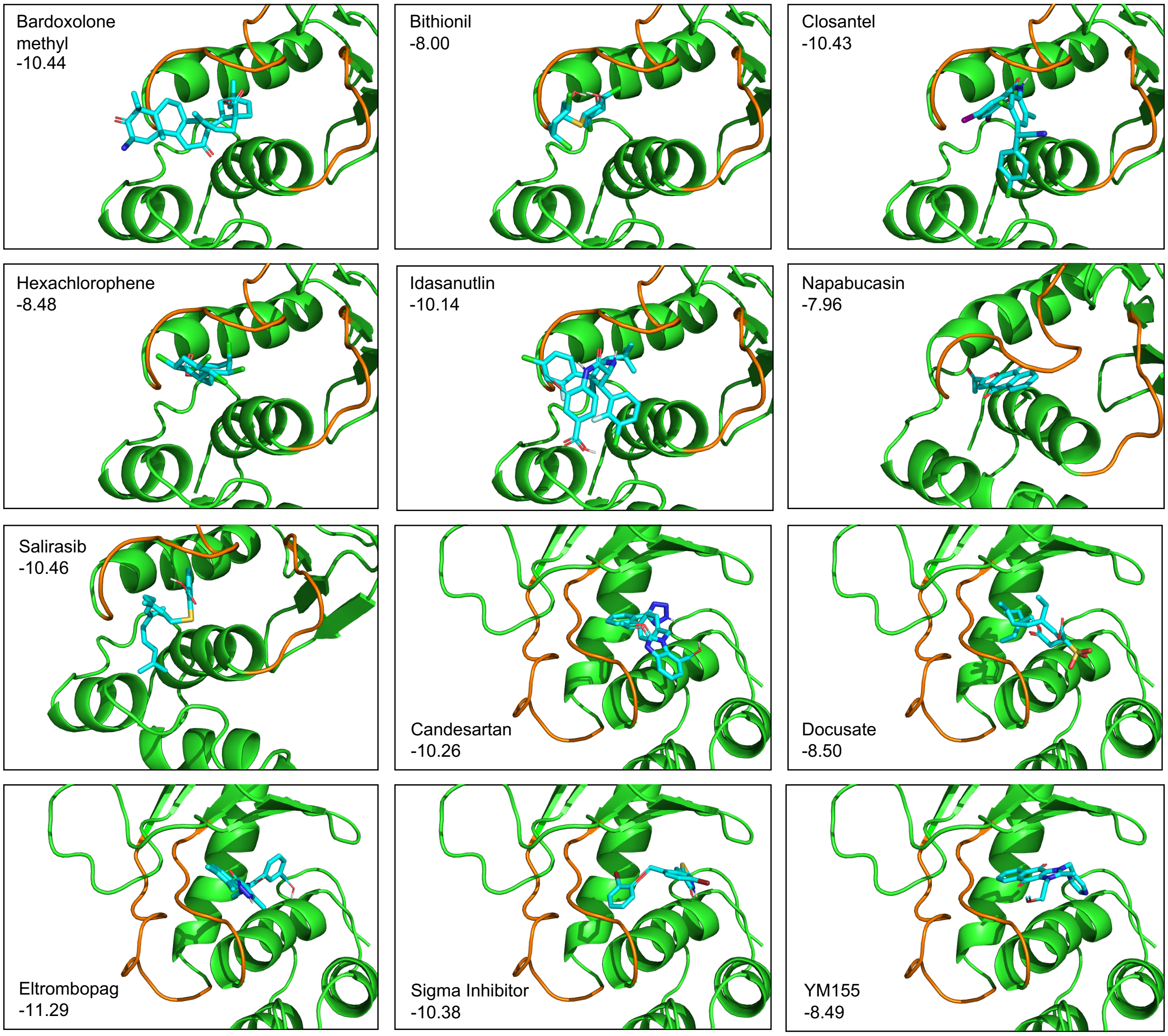


**Supplemental Figure 9. FDA-approved drugs docked onto allosteric sites of PRL-3 in closed conformation.** The binding energy for each drug is shown.

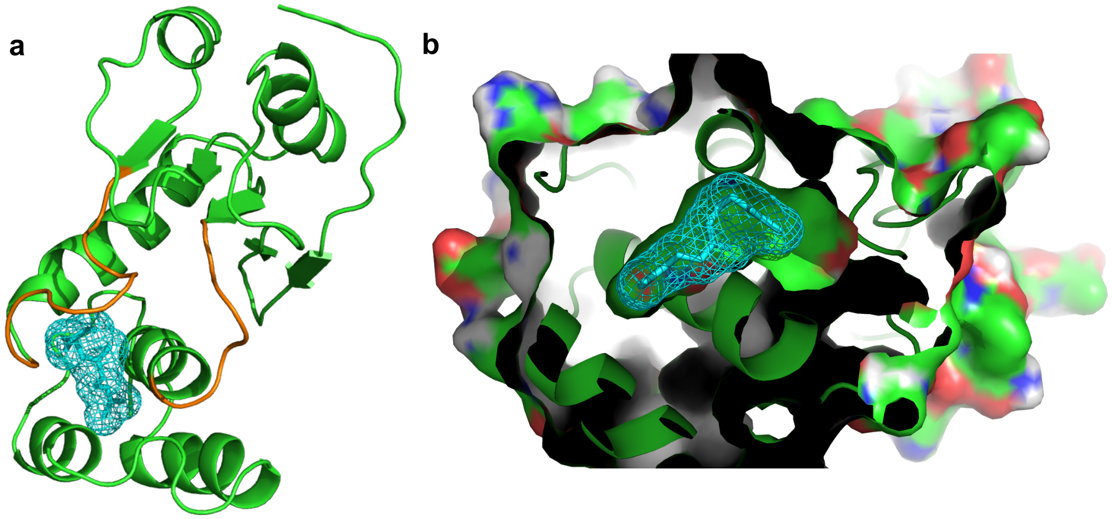

**Supplemental Figure 10. Blind docking identifies Site 1 as a potential binding site for a farnesyl tail.**
